# Privacy-preserving harmonization via distributed ComBat

**DOI:** 10.1101/2021.07.30.454516

**Authors:** Andrew A. Chen, Chongliang Luo, Yong Chen, Russell T. Shinohara, Haochang Shou, the Alzheimer’s Disease Neuroimaging Initiative

## Abstract

Challenges in clinical data sharing and the need to protect data privacy have led to the development and popularization of methods that do not require directly transferring patient data. In neuroimaging, integration of data across multiple institutions also introduces unwanted biases driven by scanner differences. These scanner effects have been shown by several research groups to severely affect downstream analyses. To facilitate the need of removing scanner effects in a distributed data setting, we introduce distributed ComBat, an adaptation of a popular harmonization method for multivariate data that borrows information across features. We present our fast and simple distributed algorithm and show that it yields equivalent results using data from the Alzheimer’s Disease Neuroimaging Initiative. Our method enables harmonization while ensuring maximal privacy protection, thus facilitating a broad range of downstream analyses in functional and structural imaging studies.

## 1 Introduction

Sharing data across medical institutions enables large-scale clinical research with more generalizable and impactful results. However, directly transferring data across organizations presents a number of issues including patient privacy concerns, incompatibility of data formats, and hardware limitations. In many cases, these concerns prevent data aggregation in their complete form. This distributed data setting has motivated several adaptations of common methods that operate without the need to share original data across sites. Recent developments have included distributed clustering (İnan et al., 2007), logistic regression (Duan et al., 2020a), Cox regression (Duan et al., 2020b), principal component analysis (Al-Rubaie et al., 2017), and deep learning (Shokri & Shmatikov, 2015).

In neuroimaging, performing analyses across multiple institutions and scanners can introduce systematic measurement errors, which are often called scanner effects. These effects can be introduced by several scanner properties including scanner manufacturer, model, magnetic field strength, head coil, voxel size, acquisition parameters, and a wide range of other differences across scanners (Han et al., 2006; Kruggel et al., 2010; Reig et al., 2009; Wonderlick et al., 2009). Differences can even persist when scanners have the exact same model and manufacturer (Shinohara et al., 2017).

Distributed analysis methods generally do not account for potential scanner effects or other types of batch effects. However, these effects are important to address and can otherwise lead to spurious associations and scanner-specific data properties that are easily detected using a classifier (Fortin et al., 2018; Glocker et al., 2019).

To mitigate scanner effects, a wide range of statistical harmonization techniques have been tested in neuroimaging data. Many of these methods address scanner effects in the mean and variance of voxel intensities or derived features (Fortin et al., 2016, 2018). Among these, ComBat (Johnson et al., 2007) has become a popular harmonization method and has been tested in both structural and functional imaging (Bartlett et al., 2018; Fortin et al., 2017; Marek et al., 2019; Yu et al., 2018). However, none of these methods can be directly applied to distributed data.

To enable harmonization in distributed data, we introduce distributed ComBat (d-ComBat), a distributed algorithm for performing ComBat. We apply our algorithm to the Alzheimer’s Disease Neuroimaging Initiative (ADNI) dataset and show that our method yields identical results to applying ComBat while having the full data at a single location. Our investigation enables additional downstream distributed methods to be applied on harmonized data and fulfills the needs for running a complete distributed analysis pipeline in multi-site neuroimaging studies.

## 2 Methods

### 2.1 Distributed ComBat

ComBat (Fortin et al., 2017, 2018; Johnson et al., 2007) seeks to remove scanner effects in the mean and variance of neuroimaging data in an empirical Bayes framework. To handle the distributed data setting, we propose d-ComBat as an algorithm that yields adjusted data identical to the original ComBat method. Let ***y**_ij_* = (*y_ij_*_1_, *y_ij_*_2_*, …, y_ijv_*)^*T*^, *i* = 1, 2*, …, K*, *j* = 1, 2, …, *n_i_* denote the *v*-dimensional vectors of observed data where *i* indexes scanner, *j* indexes subjects within scanners, *n_i_* is the number of subjects acquired on scanner *i*, and *V* is the number of features. For simplicity, we assume each site uses a different scanner and the data are collected from *K* sites. However, our algorithm could be easily extended to allow varying number of scanners per site. Our goal is to harmonize the data from these 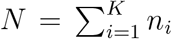 subjects across the *K* scanners without pooling data at a single processing site. ComBat assumes that the *V* features *v* = 1, 2, …, *V* follow

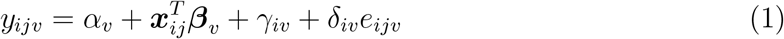

where *α_v_* is the intercept, ***x**_ij_* is the vector of covariates, ***β**_v_* is the vector of regression coefficients, *γ_iv_* is the mean scanner effect, and *δ_iv_* is the variance scanner effect. The errors *e_ijv_* are assumed to follow 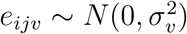.

The original ComBat contains two steps. The first is to standardize the original features by removing the covariate effects and scaling each residuals by its total variance. The second step involves estimating the scanner effects *γ* and *δ* using an empirical Bayes framework and removing them from the original data. We propose a distributed algorithm for each of the two steps in the next two sections.

#### Standardization

The original implementation of ComBat first standardizes the mean and variance of data across scanners via feature-wise least-squares estimation. The standardized data are calculated as

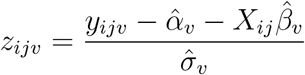

However, in the distributed setting we do not have direct access to the entire dataset and cannot directly compute estimates for the intercepts *α_v_*, regression coefficients ***β**_v_*, scanner-specific mean shifts *γ_iv_* or population standard deviations *σ_v_* for each feature. To address this problem, we propose an estimation procedure that only requires computation and transmission of deidentified summary statistics between distributed sites and a central location. As in the original ComBat methodology, estimation is performed under the constraint 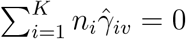 to ensure identifiability.

For each feature, define 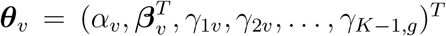. Then we can rewrite the data across all *N* subjects 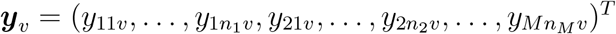 as ***y**_v_* = *W **θ*** + *e_v_* where

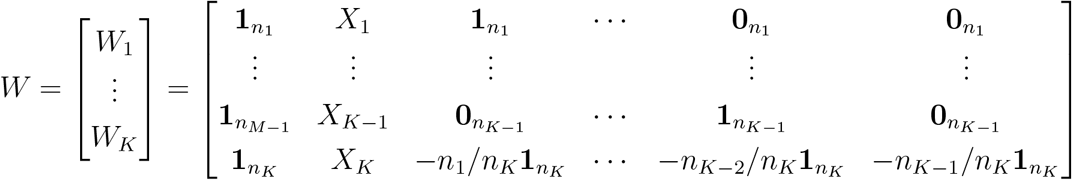

The ordinary least square estimate can be obtained via 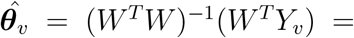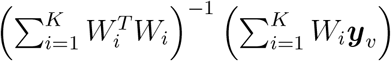. By decomposing the estimation into site-specific summary statistics 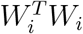 and 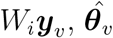 can be obtained by computing these summary statistics and sending them to a central location. Construction of *W_i_* and calculation of these summary statistics are simple for *i* = 1, 2,…, *K*−1 since they are just the usual design matrices *X_i_* concatenated with an intercept column and scanner-specific columns of ones. To standardize the variance of the data, the marginal variance is estimated as 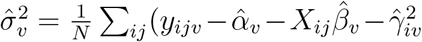 *v* = 1, 2, …, *p*, which is decomposable by site.

#### Empirical Bayes adjustment

The key step in ComBat involves use of empirical Bayes estimates of site-specific location and scale parameters to remove site effects while pooling information across features. ComBat assumes that the prior distributions 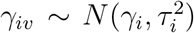 and 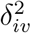 ~ Inverse Gamma(*λ_i_, ν_i_*) where hyperparameter estimates 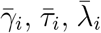, and 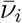 are obtained via method of moments. ComBat then finds the conditional posterior means 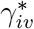 and 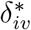, computed iteratively through

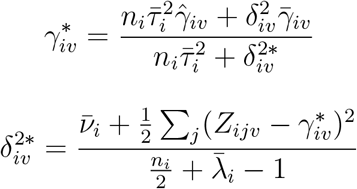

Each site’s mean and variance parameter estimates are computed from data within that site and so this step is distributed by its nature. The ComBat-adjusted data is then obtained within each site via

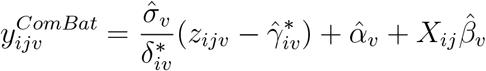

#### Algorithm

In the distributed setting, ComBat only requires two back-and-forth communications between sites and a central location for estimation of the standardization parameters. We propose the d-ComBat algorithm and illustrate our method in Fig. 1

1. Initiation - broadcast from central site: The central analysis site chooses identification numbers for each scanner and communicates these to each location.
2. Local computation at collaborative sites for mean parameters.
  a. Each site locally computes scanner-specific summary statistics 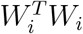and 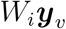 to the central site (Fig. 1a).
  b. These summary statistics are then sent back to the central site.
3. Aggregation at central site and broadcast.
  a. From the scanner-specific summary statistics, the central site computes 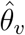.
  b. The central site then sends 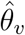 to each location (Fig. 1a).
4. Distributed data harmonizations.
  a. To obtain the global variance estimate, each site transfers 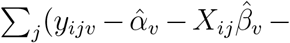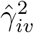 to the central location, which then sends back 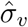 (Fig. 1b).
  b. The remaining ComBat steps are performed within each site to obtain 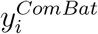 at every location (Fig. 1c).

**Figure 1:**
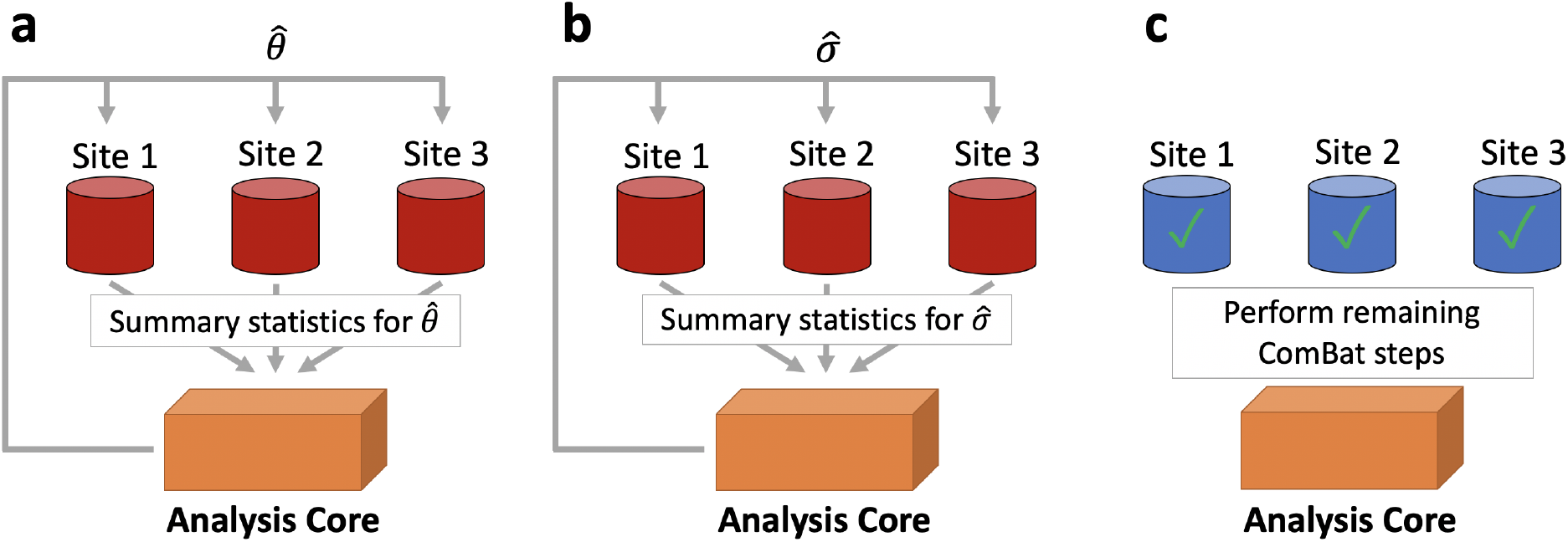
Distributed ComBat illustration. The procedure to perform distributed ComBat harmonization is outlined as follows. **a**, Each site sends its deidentified summary statistics to a central site for estimation of regression coefficients which are then passed back to the sites. **b**, Each site sends summary statistics to a central site for estimation of the population variance which is then passed back to the sites. **c**, The sites can then use the global regression coefficients and variance estimates to perform the remaining ComBat steps and obtain harmonized data.

### 2.2 ADNI data analysis

Data for our primary analysis are obtained from ADNI (http://adni.loni.usc.edu/ and processed using the ANTs longitudinal single-subject template pipeline (Tustison et al., 2019) with code available on GitHub (https://github.com/ntustison/CrossLong). All participants in the ADNI study gave informed consent and institutional review boards approved the study at all contributing institutions.

First, we obtain raw T1-weighted images from the ADNI-1 database, which were acquired using MPRAGE for Siemens and Philips scanners and a works-in-progress version of MPRAGE on GE scanners (Jack et al., 2010). For each subject, we estimate a template from all the image timepoints. Each normalized timepoint image undergoes rigid spatial normalization to this single-subject template followed by processing via a single image cortical thickness pipeline consisting of brain extraction (Avants et al., 2010), denoising (Manjón et al., 2010), N4 bias correction (Tustison et al., 2010), Atropos n-tissue segmentation (Avants et al., 2011), and registration-based cortical thickness estimation (Das et al., 2009). We include the 62 cortical thickness values from the baseline scans in our primary dataset.

We then identified scanner based on information contained within the Digital Imaging and Communications in Medicine (DICOM) headers for each scan. We consider subjects to be acquired on the same scanner if they share the scanner site, scanner manufacturer, scanner model, head coil, and magnetic field strength. In total, this definition yields 142 distinct scanners of which 78 had less than three subjects and were removed from analyses. The final sample consists of 505 subjects across 64 scanners, with 213 subjects imaged on scanners manufactured by Siemens, 70 by Philips, and 222 by GE. These 64 scanners are divided across 53 distinct ADNI sites. The sample has a mean age of 75.3 (SD 6.70) and includes 278 (55%) males, 115 (22.8%) Alzheimer’s disease (AD) patients, 239 (47.3%) late mild cognitive impairment (LMCI), and 151 (29.9%) cognitively normal (CN) individuals.

### 2.3 Comparison with ComBat

We conduct an experiment to compare d-ComBat and ComBat applied on the full data available at a single location. To emulate a distributed data setting, we treat each of the 53 ADNI sites as separate locations and only enable sharing of summary statistics with a central location. We then apply d-ComBat to this data while including age, sex, and disease status as covariates. For the reference ComBat-adjusted data, we apply ComBat including the same covariates while all of the data is housed at a single site.

We also compare these two ComBat outputs by comparing their parameter estimates, harmonized output data, and run time. Parameter estimates are compared through the maximum difference between the two sets of estimates. We then compare the harmonized data within each site and report the maximum error across all sites. For run time, we compare the ComBat run time with the time elapsed across all d-ComBat steps, including calculations at the central location.

## 3 Results

We ran d-ComBat and ComBat in R on a laptop computer running macOS Catalina version 10.15.7 with a 2.3 GHz 8-Core Intel Core i9 processor. D-ComBat ran in 387 milliseconds across all sites and steps versus ComBat which took 40 milliseconds. The average run time within each site was 7.04 milliseconds and the central site took 6 milliseconds to compute the necessary estimates.

Fig. 2 compare the empirical Bayes parameter estimates and regression coefficients obtained from each method, showing no visible differences across all parameters. The maximum percent differences between estimates were 4.17 × 10^−10^ for location parameters, 1.72 × 10^−13^ for scale parameters, and 1.19 × 10^−11^ for regression coefficients.

**Figure 2:**
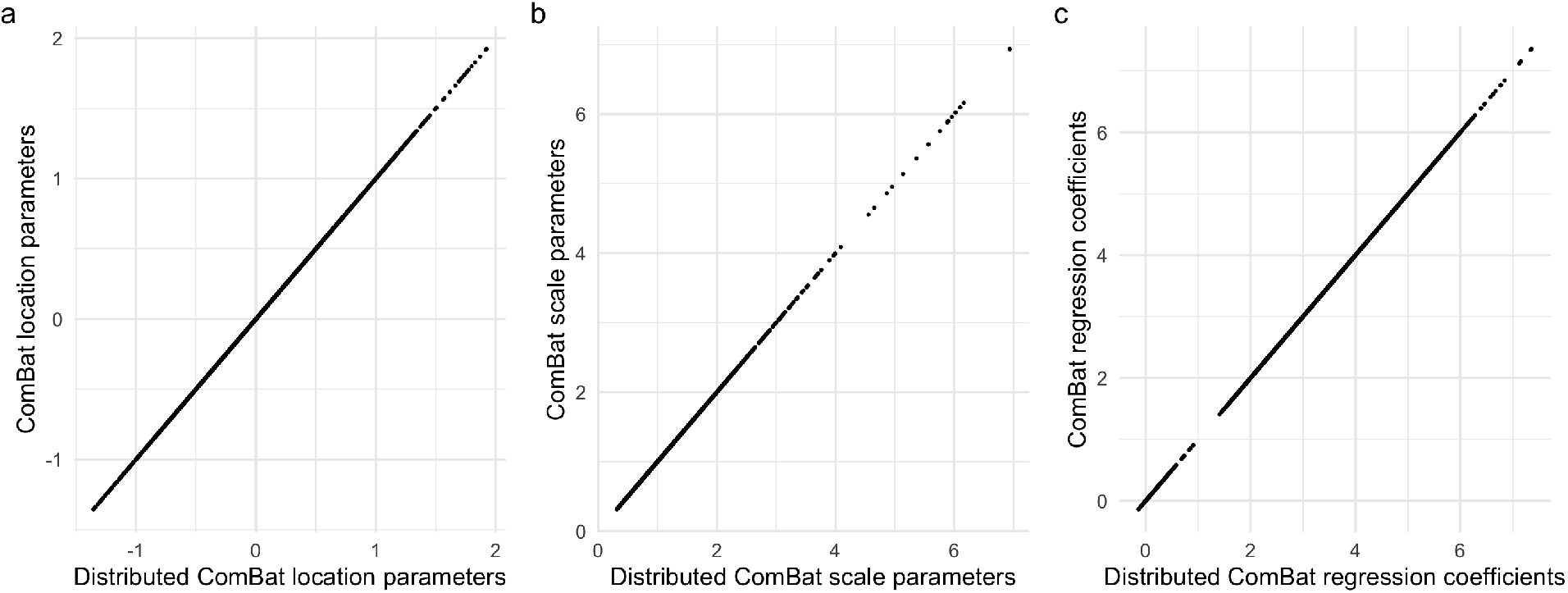
Distributed ComBat parameter estimates. Scatter plots compare parameter estimates from distributed ComBat versus those obtained from ComBat with all data at one location. **a** and **b** show empirical Bayes point estimates for location and scale respectively. **c** displays the regression coefficients obtained from each method.

The harmonized data were identical between the two methods. We found that the maximum percent difference between any two data points across the 53 locations was 2.75×10^−13^.

## 4 Discussion

Challenges in data sharing across institutions have inspired distributed algorithms for statistical analysis and machine learning. We contribute to this growing base of methods by introducing distributed ComBat for harmonization of data housed in clinical sites. To the best of our knowledge, this is the first harmonization method adapted for this setting. Compared to ComBat, we demonstrate that d-ComBat yields identical parameter estimates and harmonized output data.

Unlike ComBat, d-ComBat requires two round of communications with a central location, which requires coordination and sharing of deidentified summary statistics between sites. These additional steps result in greater total run time across all sites, but very short run times at each site. In practice, the execution time of d-ComBat will also depend on the transfer speed of summary statistics to the central location and the speed of individuals running the code at each site. The total time to run d-ComBat is likely greater than running ComBat while having data at a single location, but this additional time is expected given the complexities of a distributed data setting. Further investigation into approximating the standardization step in one communication step could greatly improve the ease of using d-ComBat.

For distributed Combat, only aggregated statistics are communicated, and the re-identification risk for the patients is expected to be low. In the future, we plan to formally quantify the re-identification risk rigorously, and enhance our algorithms via techniques including differential privacy (Dwork & Roth, 2014; Dwork et al., 2016; Wasserman & Zhou, 2010). Future studies could also adapt other harmonization methods for distributed data, including extensions of ComBat for longitudinal data (Beer et al., 2020), nonlinear associations (Pomponio et al., 2020), and covariance effects (Chen et al., 2019).

## Notes

### Competing Interest Statement

The authors have declared no competing interest.

## References

Al-Rubaie, M., Wu, P., Chang, J. M. & Kung, S. (2017). Privacy-preserving PCA on horizontally-partitioned data. In 2017 IEEE Conference on Dependable and Secure Computing.

Avants, B., Klein, A., Tustison, N., Woo, J. & Gee, J. C. (2010). Evaluation of open-access, automated brain extraction methods on multi-site multi-disorder data. In 16th Annual Meeting for the Organization of Human Brain Mapping.

Avants, B. B., Tustison, N. J., Wu, J., Cook, P. A. & Gee, J. C. (2011). An open source multivariate framework for n-tissue segmentation with evaluation on public data. Neuroinformatics 9, 381–400.

Bartlett, E. A., DeLorenzo, C., Sharma, P., Yang, J., Zhang, M., Petkova, E., Weissman, M., McGrath, P. J., Fava, M., Ogden, R. T., Kurian, B. T., Malchow, A., Cooper, C. M., Trombello, J. M., McInnis, M., Adams, P., Oquendo, M. A., Pizzagalli, D. A., Trivedi, M. & Parsey, R. V. (2018). Pretreatment and early-treatment cortical thickness is associated with SSRI treatment response in major depressive disorder. Neuropsychopharmacology 43, 2221–2230.

Beer, J. C., Tustison, N. J., Cook, P. A., Davatzikos, C., Sheline, Y. I., Shinohara, R. T. & Linn, K. A. (2020). Longitudinal ComBat: A method for harmonizing longitudinal multi-scanner imaging data. NeuroImage 220, 117129.

Chen, A. A., Beer, J. C., Tustison, N. J., Cook, P. A., Shinohara, R. T. & Shou, H. (2019). Removal of Scanner Effects in Covariance Improves Multivariate Pattern Analysis in Neuroimaging Data. bioRxiv, 858415.

Das, S. R., Avants, B. B., Grossman, M. & Gee, J. C. (2009). Registration based cortical thickness measurement. NeuroImage 45, 867–879.

Duan, R., Boland, M. R., Liu, Z., Liu, Y., Chang, H. H., Xu, H., Chu, H., Schmid, C. H., Forrest, C. B., Holmes, J. H., Schuemie, M. J., Berlin, J. A., Moore, J. H. & Chen, Y. (2020a). Learning from electronic health records across multiple sites: A communication-efficient and privacy-preserving distributed algorithm. Journal of the American Medical Informatics Association 27, 376–385.

Duan, R., Luo, C., Schuemie, M. J., Tong, J., Liang, C. J., Chang, H. H., Boland, M. R., Bian, J., Xu, H., Holmes, J. H., Forrest, C. B., Morton, S. C., Berlin, J. A., Moore, J. H., Mahoney, K. B. & Chen, Y. (2020b). Learning from local to global: An efficient distributed algorithm for modeling time-to-event data. Journal of the American Medical Informatics Association 27, 1028–1036.

Dwork, C., McSherry, F., Nissim, K. & Smith, A. (2016). Calibrating Noise to Sensitivity in Private Data Analysis. Journal of Privacy and Confidentiality 7, 17–51.

Dwork, C. & Roth, A. (2014). The Algorithmic Foundations of Differential Privacy. Foundations and Trends® in Theoretical Computer Science 9, 211–407.

Fortin, J.-P., Cullen, N., Sheline, Y. I., Taylor, W. D., Aselcioglu, I., Cook, P. A., Adams, P., Cooper, C., Fava, M., McGrath, P. J., McInnis, M., Phillips, M. L., Trivedi, M. H., Weissman, M. M. & Shinohara, R. T. (2018). Harmonization of cortical thickness measurements across scanners and sites. NeuroImage 167, 104–120.

Fortin, J.-P., Parker, D., Tunç, B., Watanabe, T., Elliott, M. A., Ruparel, K., Roalf, D. R., Satterthwaite, T. D., Gur, R. C., Gur, R. E., Schultz, R. T., Verma, R. & Shinohara, R. T. (2017). Harmonization of multi-site diffusion tensor imaging data. NeuroImage 161, 149–170.

Fortin, J.-P., Sweeney, E. M., Muschelli, J., Crainiceanu, C. M. & Shinohara, R. T. (2016). Removing inter-subject technical variability in magnetic resonance imaging studies. NeuroImage 132, 198–212.

Glocker, B., Robinson, R., Castro, D. C., Dou, Q. & Konukoglu, E. (2019). Machine Learning with Multi-Site Imaging Data: An Empirical Study on the Impact of Scanner Effects. arXiv:1910.04597 [cs, eess, q-bio].

Han, X., Jovicich, J., Salat, D., van der Kouwe, A., Quinn, B., Czanner, S., Busa, E., Pacheco, J., Albert, M., Killiany, R., Maguire, P., Rosas, D., Makris, N., Dale, A., Dickerson, B. & Fischl, B. (2006). Reliability of MRI-derived measurements of human cerebral cortical thickness: The effects of field strength, scanner upgrade and manufacturer. NeuroImage 32, 180–194.

İnan, A., Kaya, S. V., Saygın, Y., Savaş, E., Hintoğlu, A. A. & Levi, A. (2007). Privacy preserving clustering on horizontally partitioned data. Data & Knowledge Engineering 63, 646–666.

Jack, C. R., Bernstein, M. A., Borowski, B. J., Gunter, J. L., Fox, N. C., Thompson, P. M., Schuff, N., Krueger, G., Killiany, R. J., DeCarli, C. S., Dale, A. M. & Weiner, M. W. (2010). Update on the MRI Core of the Alzheimer’s Disease Neuroimaging Initiative. Alzheimer’s & dementia : the journal of the Alzheimer’s Association 6, 212–220.

Johnson, W. E., Li, C. & Rabinovic, A. (2007). Adjusting batch effects in microarray expression data using empirical Bayes methods. Biostatistics 8, 118–127.

Kruggel, F., Turner, J., Muftuler, L. T. & Alzheimer’s Disease Neuroimaging Initiative (2010). Impact of scanner hardware and imaging protocol on image quality and compartment volume precision in the ADNI cohort. NeuroImage 49, 2123–2133.

Manjón, J. V., Coupé, P., Martí-Bonmatí, L., Collins, D. L. & Robles, M. (2010). Adaptive non-local means denoising of MR images with spatially varying noise levels. Journal of magnetic resonance imaging: JMRI 31, 192–203.

Marek, S., Tervo-Clemmens, B., Nielsen, A. N., Wheelock, M. D., Miller, R. L., Laumann, T. O., Earl, E., Foran, W. W., Cordova, M., Doyle, O., Perrone, A., Miranda-Dominguez, O., Feczko, E., Sturgeon, D., Graham, A., Hermosillo, R., Snider, K., Galassi, A., Nagel, B. J., Ewing, S. W. F., Eggebrecht, A. T., Garavan, H., Dale, A. M., Greene, D. J., Barch, D. M., Fair, D. A., Luna, B. & Dosenbach, N. U. F. (2019). Identifying reproducible individual differences in childhood functional brain networks: An ABCD study. Developmental Cognitive Neuroscience 40, 100706.

Pomponio, R., Erus, G., Habes, M., Doshi, J., Srinivasan, D., Mamourian, E., Bashyam, V., Nasrallah, I. M., Satterthwaite, T. D., Fan, Y., Launer, L. J., Masters, C. L., Maruff, P., Zhuo, C., Völzke, H., Johnson, S. C., Fripp, J., Koutsouleris, N., Wolf, D. H., Gur, R., Gur, R., Morris, J., Albert, M. S., Grabe, H. J., Resnick, S. M., Bryan, R. N., Wolk, D. A., Shinohara, R. T., Shou, H. & Davatzikos, C. (2020). Harmonization of large MRI datasets for the analysis of brain imaging patterns throughout the lifespan. NeuroImage 208, 116450.

Reig, S., Sánchez-González, J., Arango, C., Castro, J., González-Pinto, A., Ortuño, F., Crespo-Facorro, B., Bargalló, N. & Desco, M. (2009). Assessment of the increase in variability when combining volumetric data from different scanners. Human Brain Mapping 30, 355–368.

Shinohara, R. T., Oh, J., Nair, G., Calabresi, P. A., Davatzikos, C., Doshi, J., Henry, R. G., Kim, G., Linn, K. A., Papinutto, N., Pelletier, D., Pham, D. L., Reich, D. S., Rooney, W., Roy, S., Stern, W., Tummala, S., Yousuf, F., Zhu, A., Sicotte, N. L., Bakshi, R. & Cooperative, t. N. (2017). Volumetric Analysis from a Harmonized Multisite Brain MRI Study of a Single Subject with Multiple Sclerosis. American Journal of Neuroradiology 38, 1501–1509.

Shokri, R. & Shmatikov, V. (2015). Privacy-Preserving Deep Learning. In Proceedings of the 22nd ACM SIGSAC Conference on Computer and Communications Security, CCS’15. New York, NY, USA: Association for Computing Machinery.

Tustison, N. J., Avants, B. B., Cook, P. A., Zheng, Y., Egan, A., Yushkevich, P. A. & Gee, J. C. (2010). N4ITK: Improved N3 Bias Correction. IEEE Transactions on Medical Imaging 29, 1310–1320.

Tustison, N. J., Holbrook, A. J., Avants, B. B., Roberts, J. M., Cook, P. A., Reagh, Z. M., Duda, J. T., Stone, J. R., Gillen, D. L., Yassa, M. A. & Initiative, f. t. A. D. N. (2019). Longitudinal Mapping of Cortical Thickness Measurements: An Alzheimer’s Disease Neuroimaging Initiative-Based Evaluation Study. Journal of Alzheimer’s Disease 71, 165–183.

Wasserman, L. & Zhou, S. (2010). A Statistical Framework for Differential Privacy. Journal of the American Statistical Association 105, 375–389.

Wonderlick, J., Ziegler, D., Hosseini-Varnamkhasti, P., Locascio, J., Bakkour, A., van der Kouwe, A., Triantafyllou, C., Corkin, S. & Dickerson, B. (2009). Reliability of MRI-derived cortical and subcortical morphometric measures: Effects of pulse sequence, voxel geometry, and parallel imaging. NeuroImage 44, 1324–1333.

Yu, M., Linn, K. A., Cook, P. A., Phillips, M. L., McInnis, M., Fava, M., Trivedi, M. H., Weissman, M. M., Shinohara, R. T. & Sheline, Y. I. (2018). Statistical harmonization corrects site effects in functional connectivity measurements from multi-site fMRI data. Human Brain Mapping 39, 4213–4227.

